# Sequential Specification of Oligodendrocyte Lineage Cells by Distinct Levels of Hedgehog and Notch Signaling

**DOI:** 10.1101/377671

**Authors:** Andrew M. Ravanelli, Christina A. Kearns, Rani K. Powers, Yuying Wang, Jacob H. Hines, Maranda J. Donaldson, Bruce Appel

**Author notes:** Current addresses: A.M.R., MilliporeSigma St. Louis, Missouri, 2909 Laclede Ave, USA Y.W., BGI, Shenzhen, China J.H.H., Biology Department, Winona State University, Winona MN 55987. Lead contact: Bruce Appel.

## Abstract

During development of the central nervous system oligodendrocyte precursor cells (OPCs) give rise to both myelinating oligodendrocytes and NG2 glia, which are the most proliferative cells in the adult mammalian brain. NG2 glia retain characteristics of OPCs, and some NG2 glia produce oligodendrocytes, but many others persist throughout adulthood. Why some OPCs differentiate as oligodendrocytes during development whereas others persist as OPCs and acquire characteristics of NG2 glia is not known. Using zebrafish spinal cord as a model, we found that OPCs that differentiate rapidly as oligodendrocytes and others that remain as OPCs arise in sequential waves from distinct neural progenitors. Additionally, oligodendrocyte and persistent OPC fates are specified during a defined critical period by small differences in Shh signaling and Notch activity, which modulates Shh signaling response. Thus, our data indicate that OPCs fated to produce oligodendrocytes or remain as OPCs during development are specified as distinct cell types, raising the possibility that the myelinating potential of OPCs is set by graded Shh signaling activity.

## INTRODUCTION

Myelin, a specialized, proteolipid-rich membrane, tightly wraps axons to enhance their conduction of electrical impulses and provide them with metabolic support (Simons and Nave, 2015). In the central nervous system of vertebrate animals, myelin is produced by glial cells called oligodendrocytes. Following neurogenesis, subpopulations of neural progenitors give rise to oligodendrocyte precursor cells (OPCs), which divide and migrate to populate the neural tube. During postnatal stages, many OPCs stop dividing and differentiate as oligodendrocytes. However, other OPCs persist into adulthood (Dawson et al., 2003). These cells often are called NG2 glia because they express the chondroitin sulfate proteoglycan NG2, encoded by *CSPG4*, and, although they can differentiate as oligodendrocytes, they have properties that are independent of oligodendrocyte formation (Nishiyama et al., 2016). Why some OPCs differentiate as oligodendrocytes during developmental stages whereas others remain as NG2 glia into adulthood is not known.

In embryos, most spinal cord OPCs arise from ventral pMN progenitors, which express the transcription factor Olig2. During early stages of neural tube development a ventral to dorsal gradient of Sonic Hedgehog (Shh) signaling activity specifies pMN progenitors, as well as a more ventral population called p3 and more dorsal p0, p1 and p2 progenitors. Each of these progenitor populations produce first neurons and then glial cells. In particular, pMN progenitors produce motor neurons prior to forming OPCs. The mechanisms that cause pMN progenitors to switch from motor neuron to OPC production are not known, but several lines of evidence indicate that modulation of Shh signaling, resulting in a temporal rise in Shh signaling activity, is concurrent with and necessary for OPC formation (Agius et al., 2004; Danesin et al., 2006; Fu et al., 2002; Oustah et al., 2014; Touahri et al., 2012; Zhou et al., 2001).

If modulation of Shh signaling helps specify pMN progenitors for motor neuron or OPC fates, could differences in Shh activity also determine if OPCs differentiate as oligodendrocytes during development or remain as OPCs that adopt characteristics of NG2 glia? Here we report a series of experiments designed to test this possibility, using zebrafish as a model system. Our main findings reveal two subpopulations of OPCs in the zebrafish larval spinal cord. These OPC subpopulations arise from distinct progenitors that sequentially initiate *olig2* expression, with those that become oligodendrocytes initiating *olig2* expression first and those that express *olig2* later remaining as OPCs. We also found that the developmental fate of OPCs could be modulated by small changes in Hedgehog and Notch signaling during the time of their specification. We previously presented evidence that motor neurons and OPCs arise from distinct progenitors that sequentially express *olig2*, a process that we called progenitor recruitment (Ravanelli and Appel, 2015). We now propose that progenitor recruitment also can account for sequential formation of distinct OPCs in spinal cord development.

## MATERIALS AND METHODS

### Experimental model details

Transgenic and mutant zebrafish lines, listed below in the Resources Table, were generated and maintained using the AB strain. Embryos were produced by pairwise matings of male and female adult fish. Embryos were raised at 28.5°C in E3 medium [5 mM NaCl, 0.17 mM KCl, 0.33 mM CaCl, 0.33 mM MgSO_4_, pH to 7.4 with sodium bicarbonate] and staged according to hours post-fertilization (hpf) and morphological criteria (Kimmel et al., 1995). Developmental stages used for individual experiments are noted in the text and figure legends. Sex cannot be determined at embryonic and larval stages. For pharmacological experiments, embryos were randomly assigned to control and experimental conditions. The University of Colorado Anschutz Medical Campus Institutional Animal Care and Use Committee (IACUC) approved all zebrafish studies and all experiments that were performed conformed to IACUC standards.

### Imaging

Confocal time-lapse imaging was carried out by embedding embryos in 0.6% low melting point agarose (IBI Scientific) with 6% tricaine (Sigma) and viewing the spinal cord in transverse planes using 20× (n.a. 0.8), 40× oil immersion (n.a. 1.3) immersion or 40× long-working-distance water immersion (n.a. 1.1) objectives. Embryos were maintained at 28.5°C in a heated stage chamber and z-stacks were collected at variable intervals, usually starting at 24 hpf and continuing through to 48 hpf. Images were taken using a Zeiss Axiovert 200 microscope equipped with a PerkinElmer spinning disk confocal system (PerkinElmer Improvision). Image brightness and contrast were adjusted in Volocity (PerkinElmer) or ImageJ (National Institutes of Health). For comparison purposes, the same adjustment was applied to all images. Time-lapse videos were exported from Volocity as extended *z*-projection TIFF images. We used ImageJ to rotate, crop and translate in order to correct for x-y drift. Image stacks were then exported in QuickTime (.mov) format.

To image tissue sections, larvae were fixed overnight in 4% paraformaldehyde in PBST (0.1% Tween-20), embedded in agarose, frozen and sectioned using a cryostat microtome to obtain 20 um sections. Confocal images were obtained using the microscope described above.

### Kaede photoconversion

Photoconversion of *olig2*:Kaede was performed as previously described (Ravanelli and Appel, 2015). *olig2*:Kaede transgenic embryos were illuminated with UV light from a metal halide fluorescent light source for 30-120 s following manual dechorionation, or for 10 seconds with a 405 nm laser (20% power) on a spinning disk confocal microscope. Immediately afterward, photoconversion was confirmed by observation of the presence of red fluorescence and an absence of green fluorescence. Photoconverted fish were then kept at 28.5°C in the dark until imaging.

### Pharmacological inhibitor treatments

Cyclopamine (Cat#11321, Cayman Chemical) was reconstituted in ethanol to make a 10 mM stock and stored at −20°C. LY411575 (Cat#4-0054, Stemgent) was reconstituted in DMSO to make a 10 mM stock and stored at −20°C. To test the effects of reduced Hh signaling for specification, embryos were treated from 24-72 hpf with 0.05 μM or 0.5 μM cyclopamine or an equal concentration of ethanol alone in E3 medium. To test the effects of Notch signaling manipulation, embryos were treated from 24-72 hpf with 1 μM or 5 μM LY411575 or an equal concentration of DMSO alone in E3 medium. Following treatments, embryos were washed in fresh E3 medium 2 times for 30 minutes, then incubated in fresh E3 medium until imaging. To test the effects of Hh signaling manipulation on differentiation, embryos were treated from 72-120 hpf with 10 μM cyclopamine or an equal concentration of ethanol alone in E3 medium. To test the effects of Notch signaling manipulation on differentiation, embryos were treated from 72-120 hpf with 10 μM or 25 μM LY411575 or an equal concentration of DMSO alone in E3 medium. Larvae were immediately imaged at 5 dpf following treatment. To test the effects of Hh signaling manipulation on Notch signaling, *her4.3*:dRFP embryos were treated from 24-48 hpf with 5 μM cyclopamine or an equal concentration of ethanol alone in E3 medium. To test the effects of Notch signaling manipulation Hh signaling, *Tg(nkx2.2a:EGFP-CaaX)* embryos were treated from 24-48 hpf with 1 μM LY411575 or an equal concentration of DMSO alone in E3 medium. Embryos were immediately imaged at 48 hpf following treatment.

The numbers of experimental trials and embryos or larvae analyzed for each condition are noted in the text and figure legends.

### Heat-shock promoter activation

To activate the *hsp70l:Shh-GFP* transgene, embryos were manually dechorionated and placed in 2 mL of E3 medium in a 15 mL conical tube. The tube was placed in a 38°C circulating water bath for 30 minutes. Embryos were recovered in fresh 28.5°C E3 medium and cultured at 28.5°C until the time of imaging. For quantification of cell numbers, embryos were sorted at 6 hours post heat-shock using a stereomicroscope equipped with fluorescence optics to select those with high intensity Shh-GFP expression. For cell counts of activated Shh with Notch inhibitor, *hsp70l:Shh-GFP* embryos were heat-shocked at 24 hpf, sorted for Shh-GFP fluorescence and treated with 0.5 μM LY411575 or DMSO in E3 medium from 24-72 hpf, then imaged at 5 dpf. For qRT-PCR analysis of this condition, *hsp70l:Shh-GFP* embryos were heat-shocked at 24 hpf, sorted for Shh-GFP fluorescence and treated with 0.5 μM LY411575 or DMSO in E3 medium from 24-48 hpf, then RNA was harvested from embryos at 48 hpf.

The numbers of experimental trials and embryos or larvae analyzed for each condition are noted in the text and figure legends.

### Semi-quantitative RT-PCR

Total mRNA was harvested from 20 pooled embryos for each control or experimental condition. RNA isolation was performed using Trizol extraction. cDNA was synthesized using an iScript Reverse Transcription kit (Biorad, 170-8840). Real-time RT-PCR was performed in triplicate for each cDNA sample using an Applied Biosystems StepOne Plus machine and software version 2.1. Taqman Gene Expression Assays (Applied Biosystems) were used to detect *her4* (Dr03160688_g1), *ptch2* (Dr03118687_m1), *gli1* (Dr03093669_m1) and *rpl13a* (Dr03101115_g1) as the endogenous control.

### In situ RNA hybridization

Antisense, digoxygenin-labeled RNA probes were synthesized using linearized plasmids as templates. *pBlueScript:ptch2* (Concordet et al., 1996) (ZFIN ID: ZDB-GENE-980526-44) was a gift from Phil Ingham. *pCRScript:nkx2.2a* (Barth and Wilson, 1995) (ZFIN ID: ZDB-Gene-980526-403) was a gift from Steve Wilson. Hybridization and detection followed standard procedures (Hauptmann and Gerster, 2000).

### Immunohistochemistry

Larvae were fixed using 4% paraformaldehyde, embedded, frozen and sectioned in 15 μm increments using a cryostat microtome. We used rabbit anti-Sox10 antibody (Park et al., 2005a) at 1:750 dilution followed by Alexa Fluor 647 goat anti-rabbit IgG antibody (1:500, Jackson Laboratories). *olig2*:EGFP was detected using anti-GFP antibody (1:1000, Living Colors anti-GFP, Clontech #632380). Images were captured using a Zeiss 880 Laser Scanning Microscope.

### Transgenic reporter construction

Zebrafish *mbpa* regulatory DNA was amplified by PCR from genomic DNA using the primers 5’-GTCGACCAGATGCTGAGATGTGACTACTGCAAATGA-3’ and 5’-GGATCCGTTGATCTGTTCAGTGGTCTACAGTCTGGA-3’, which were described previously (Jung et al., 2010). The plasmid *pEXPR-2.6mbp:tagRFPt* was created using the Gateway (Invitrogen) Tol2 system. The transgenic line *Tg(−2.6mbp:tagRFPt)*^*co25*^ was created by coinjecting plasmid with mRNA encoding Tol2 transposase into newly fertilized eggs. These were raised to adulthood and screened for germline transmission by outcrossing to non-transgenic fish.

To create *Tg(cspg4:mCherry)*^*co28*^, we followed a published method for bacterial artificial chromosome (BAC) engineering (Bussmann and Schulte-Merker, 2011a). pCR8GW-iTol2-AmpR and pCS2-mCherry-KanR clones were obtained from the authors. A *cspg4* Bac clone (CH211-9J11) was purchased from BACPAC Resources (http://bacpacresources.org/zebrafish211.htm). The following primers were used to PCR amplify a mCherry-Kanamycin cassette from pCS2-mCherry-KanR with *cspg4* homology arms: cspg4_HA1_mcherry_fw: CTCCAGGTCCCAAAGTGGCCACAGAGACTCAGAGACTCGGACTAAAGTGGCACCA TGGTGAGCAAGGGCGAAGAG cspg4_HA2_kan_rev: AGGTATAGGAGTGCCAGGAAGAGGGCAGACAGGAGCGGACACGGGGCTCTTCAGAAGAACTCGTCAAGAAGGCG. The recombined BAC clone was miniprepped and injected into newly fertilized zebrafish eggs. Germline transmission was identified by mating adult fish to wild-type fish and examining embryos for reporter gene expression using a stereomicroscope equipped with fluorescence optics.

### Cell Dissociation and FACS

7dpf *Tg(olig2:EGFP;cspg4:mCherry)* and *Tg(olig2:EGFP;mbpa:tagRFPt)* euthanized larvae were collected in 1.7 ml microcentrifuge tubes and lysed in 1xDBS (Sigma), cell dissociation media (Accumax) and DNaseI (Roche) at 37° C for 1 hour. After being transferred to ice, 1×DPBS and DNAseI were added to the solution and filtered through 70um nylon mesh strainer (Fisher) into a conical tube. The solution was transferred to a 1.7 ml tube, centrifuged and after removing supernatant, the pellet was resuspended in sorting buffer and vital dye DAPI (Sigma) was added to mark dead cells. Cells were FAC sorted to distinguish EGFP^+^, tagRFPt^+^ and mCherry^+^ cells using a MoFlo XDP100 cell sorter at the CU-SOM Cancer Center Flow Cytometry Shared Resource. Following sorting, Trizol LS (Life Techologies) was added to FACs sorted cells. 20% Chloroform was added to the solution and centrifuged at 4° C for 15 min. The supernatant was transferred to a new tube and equal parts cold isopropanol and 2 ul glycogen blue was added and incubated overnight at −20° C. The solution was centrifuged and supernatant removed. The pellet was washed with cold EtOH and resuspended in nuclease free water then heated to 55° C for 10 min and stored at −80° C.

### RNA-seq

Library construction and RNA-seq were performed by the University of Colorado Anschutz Medical Campus Microarray and Genomics Core. Following Bioanalyzer quality assessment, RNA isolated from sorted cells with high RNA integrity number (>8) was used for library preparation using the Nugen Ovation RNA Seq Library Kit. This was followed by sequencing using an Illumina HiSEQ2500 utilizing single read 50 cycle sequencing. Three libraries, constructed from independent, pooled samples, were sequenced for each cell type.

### Quantification and statistical analysis

Quantifications of cell numbers were performed by collecting confocal *z*-stacks of entire embryos and counting cells from maximally projected images. *z* images were examined to distinguish between individual cells. Quantification of *her4.3*:dRFP and *ptch2*:Kaede fluorescence intensity was performed by measuring average pixel intensity across the spinal cord in ImageJ (National Institutes of Health), using TIFF images of embryos collected at identical exposures and normalized to area. We plotted all data and performed all statistical analyses in Microsoft Excel and Prism. All data are expressed as mean ± SEM. For statistical analysis, we used Student’s two-tailed *t*-test for all data with normal distributions. Unless otherwise noted, statistical significance is indicated as follows: \**p*<0.05, \*\**p*<0.01, \*\*\**p*<0.001, \*\*\*\**p*,0.0001.

### Key Resources Table

All zebrafish lines and reagents are detailed in the Key Resources Table.

## RESULTS

### OPCs with distinct developmental fates initiate *olig2* expression at different times

The objective of our study was to understand why, during development, some OPCs differentiate as oligodendrocytes whereas others do not. In mammals, large numbers of OPCs persist into adulthood. These cells, which often are called NG2 glia (Dawson et al., 2000) or sometimes polydendrocytes (Nishiyama et al., 2009) are defined by highly branched morphologies, expression of *Sox10, Olig2, Pdgfra* and *Cspg4*, which encodes the proteoglycan Neural/glial antigen 2 (NG2), and lack of expression of myelin protein-encoding genes, such as *Mbp*. As a first step in our analysis, we set out to better define oligodendrocyte lineage cells in the spinal cord of zebrafish larvae. Similar to mammals, post-embryonic zebrafish have both oligodendrocytes and cells with the highly branched morphologies characteristic of OPCs (Figure 1A,B). To develop a marker that distinguishes OPCs from oligodendrocytes in zebrafish, we used bacterial artificial chromosome modification to create the transgene *Tg(cspg4:mCherry)*, which expresses mCherry fluorescent protein under control of cis-regulatory elements of the *cspg4* gene. After establishing stocks that transmit the transgene through the germline, we examined *cspg4:*mCherry expression in the spinal cord by comparing it to other oligodendrocyte lineage cell markers at stages encompassing larval and adult stages. The earliest we detected *cspg4*:mCherry was 5 days post fertilization (dpf), about 80 hr after pMN progenitors produce the first OPCs and approximately 2 days after oligodendrocytes begin to wrap axons and express myelin-encoding genes. From 5-20 dpf, nearly all *cspg4*:mCherry^+^ cells expressed the oligodendrocyte lineage markers Sox10, detected by immunohistochemistry, and *olig2*:EGFP (Figure 1C-E’). However, only a subset of Sox10^+^ *olig2*:EGFP^+^ cells expressed *cspg4*:mCherry. Whereas the number of *cspg4*:mCherry^+^ cells remained similar at 5, 10 and 20 dpf, the number of Sox10^+^ *olig2*:EGFP^+^ *cspg4*:mCherry^−^ cells was substantially greater at 20 dpf than at 5 and 10 dpf (Figure 1F). Adult animals also expressed *cspg4*:mCherry as a fraction of the *olig2*:EGFP^+^ population (Figure 1G,G’). We next examined expression of *cspg4*:mCherry in combination with Sox10 and *mbpa:*tagRFPt, which drives expression of red fluorescent protein in oligodendrocytes under control of *mbpa* cis-regulatory sequence. At each stage most Sox10^+^ cells expressed either *cspg4*:mCherry or *mbpa:*tagRFPt, although a small number of Sox10^+^ cells expressed both reporters (Figure 1H-K). As above, the number of *cspg4*:mCherry^+^ cells remained consistent across stages but the number of *cspg4*:mCherry^−^ *mbpa:*tagRFPt^+^ cells increased sharply between 10 and 20 dpf (Figure K). Thus, *cspg4*:mCherry and *mbpa:*tagRFPt expression mark complementary and mostly non-overlapping populations of Sox10^+^ oligodendrocyte lineage cells. To validate the specificity of the *cspg4*:mCherry reporter, we analyzed RNA-seq data obtained from transgenically marked cells that had been isolated from 7 dpf larvae. By this measure, *cspg4*:mCherry^+^ *olig2*:EGFP^+^ cells expressed *cspg4* transcripts at 50-fold greater levels than *mbpa:*tagRFPt^+^ *olig2*:EGFP^+^ cells (Figure 1L). Additionally, *cspg4*:mCherry^+^ *olig2*:EGFP^+^ cells expressed *pdgfra* at approximately 8-fold greater levels than *mbpa:*tagRFPt^+^ *olig2*:EGFP^+^ cells (Figure 1L). Together, these data indicate that zebrafish larvae produce oligodendrocyte lineage cells that differentiate as oligodendrocytes or persist as *cspg4^+^* OPCs.

**Figure 1.**
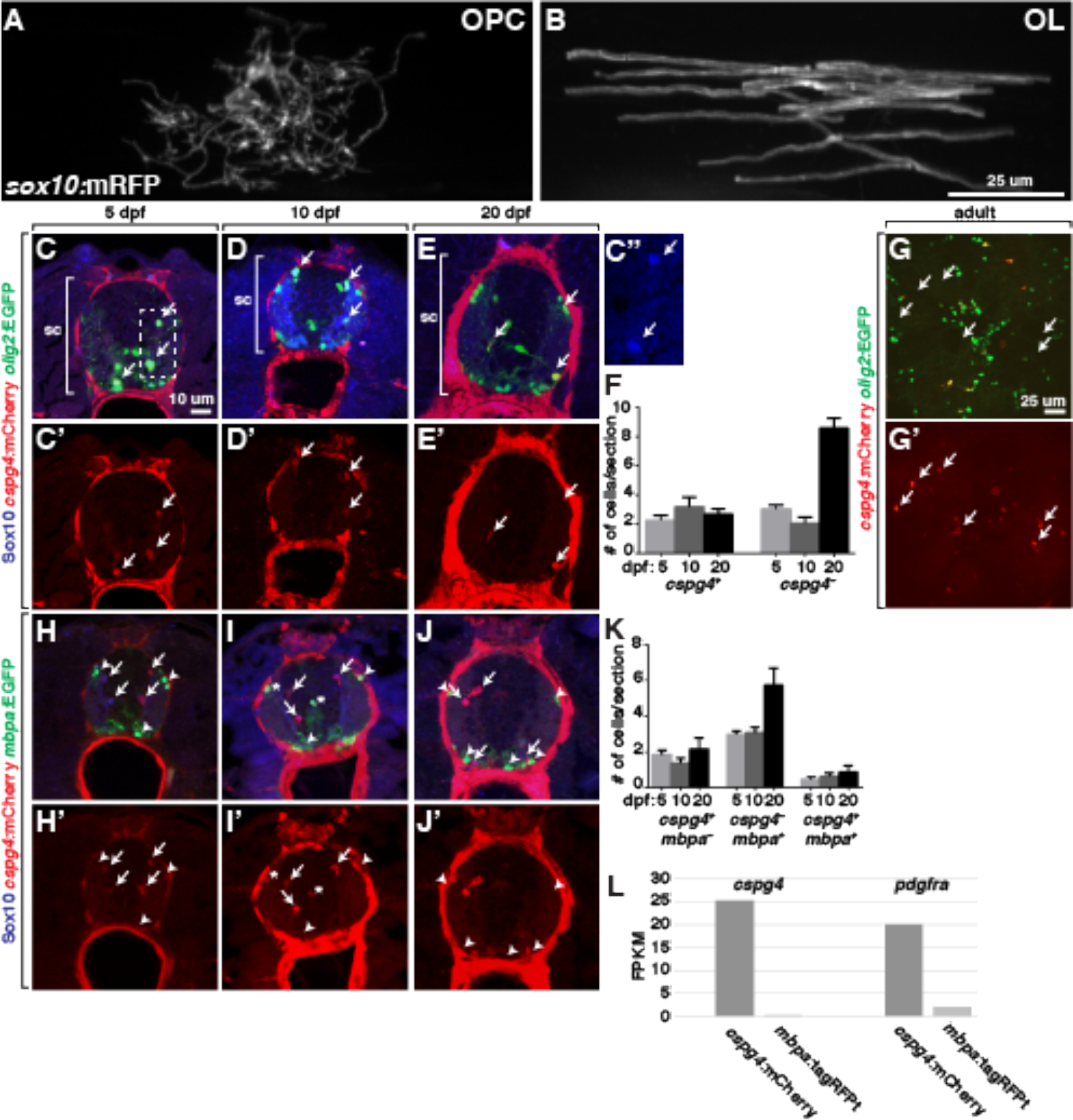
OPCs and Oligodendrocytes Occupy Larval Zebrafish Spinal Cord. (A) *sox10*:mRFP^+^ cell in a 7 dpf larvae with morphology characteristic of an OPC. (B) *sox10*:mRFP^+^ oligodendrocyte (OL) at 7 dpf. (C-E) Representative confocal images of transverse sections at the level of trunk spinal cord (sc, brackets) of 5, 10 and 20 dpf larvae showing *cspg4:*mCherry and *olig2*:EGFP expression combined with Sox10 immunohistochemistry. Examples of triple-labeled cells are indicated by arrows. C’-E’ show *cspg4:*mCherry alone. C” is an enlargement of the boxed area in panel C to show Sox10^+^ cells. (F) Number of OPCs (*cspg4:*mCherry^+^ *olig2*:EGFP^+^ Sox10^+^) and oligodendrocytes (*cspg4:*mCherry^−^ *olig2*:EGFP^+^ Sox10^+^) per transverse section for 5, 10 and 20 dpf larval stages. n=10 larvae for each stage. (G) Transverse section of adult spinal cord showing *cspg4:*mCherry and *olig2*:EGFP expression. Arrows indicate examples of double-labeled cells. G’ shows *cspg4:*mCherry alone. (H-J) Representative confocal images of transverse sections at the level of trunk spinal cord of 5, 10 and 20 dpf larvae showing *cspg4:*mCherry and *mbpa*:EGFP expression combined with Sox10 immunohistochemistry. Arrows indicate *cspg4:*mCherry^+^ *mbpa*:EGFP^−^ Sox10^+^ OPCs, arrowheads mark *cspg4:*mCherry^−^ *mbpa*:EGFP^+^ Sox10^+^ oligodendrocytes and asterisks indicate rare *cspg4:*mCherry^+^ *mbpa*:EGFP^+^ Sox10^+^ cells. (K) Number of OPCs (*cspg4:*mCherry^+^ *mbpa*:EGFP^−^ Sox10^+^) and oligodendrocytes (*cspg4:*mCherry^−^ *mbpa*:EGFP^+^ Sox10^+^ and *cspg4:*mCherry^+^ *mbpa*:EGFP^+^ Sox10^+^) per transverse section for 5, 10 and 20 dpf larval stages. n=10 larvae for each stage. (L) Expression (FPKM) of *cspg4* and *pdgfra* in *cspg4:*mCherry^+^ and *mbpa*:tagRFPt^+^ cells isolated from 7 dpf larvae.

Although *cspg4:*mCherry expression marks zebrafish spinal cord OPCs, it is evident only after the first oligodendrocytes, marked by *mbpa* expression, begin to differentiate at 3 dpf. Thus, OPCs that produce the earliest-myelinating oligodendrocytes apparently do not express *cspg4*. Are OPCs that differentiate rapidly as oligodendrocytes or persist to express *cspg4* distinct populations of cells? We previously presented evidence that the subset of OPCs that differentiate as oligodendrocytes at early larval stage express *nkx2.2a:*EGFP-CaaX (Kucenas et al., 2008b). Consistent with our observation above that most *mbpa:*tagRFPt^+^ oligodendrocytes do not express *cspg4*:mCherry, we found no evidence that *cspg4:*mCherry^+^ OPCs express *nkx2.2a:*EGFP-CaaX at 5 dpf (Figure 2A-A”). To learn if *nkx2.2a:*EGFP-CaaX expression reveals distinct subsets of OPCs during embryonic development we performed time-lapse imaging experiments using the transgenic reporter combination *Tg(olig2:DsRed2);Tg(nkx2.2a:EGFP-CaaX)*. At 53 hpf, nearly all dorsally migrated *olig2:*DsRed2^+^ cells were also *nkx2.2a*:EGFP-CaaX^+^ (41 of 42 OPCs, n=4 embryos, Figure 2B,B’), indicating that these cells were fated to differentiate as oligodendrocytes. By 60 hpf, the total number of dorsal *olig2:*DsRed2^+^ cells had increased to 68, of which 23 were *nkx2.2a*:EGFP-CaaX^−^ (Figure 2C,C’). Thus, early-migrating OPCs expressed *nkx2.2a*:EGFP-CaaX whereas later migrating OPCs did not, consistent with the possibility that distinct subpopulations of OPCs occupy the spinal cord of zebrafish embryos.

**Figure 2.**
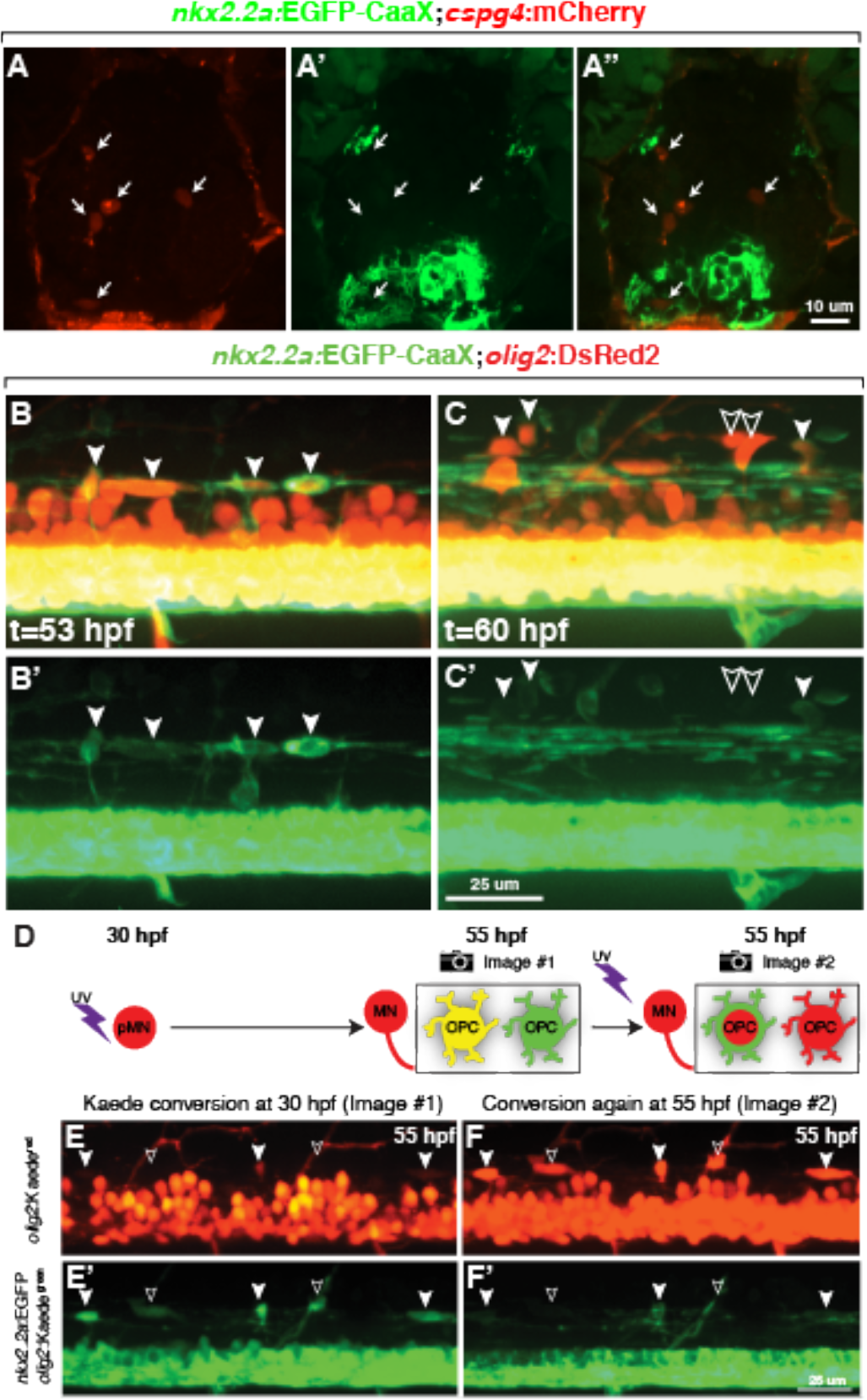
Distinct OPCs Arise Sequentially from Progenitors that Initiate *olig2* Expression at Different Times. (A-A”) Representative transverse section of 5 dpf *Tg(nkx2.2a:EGFP-CaaX);Tg(cspg4:mCherry)* larva showing red, green and combined channels. Arrows indicate *cspg4*:mCherry^+^ OPCs. (B-C’). Images captured from time-lapse videos of *Tg(olig2:DsRed2);Tg(nkx2.2a:EGFP-CaaX)* embryos at the level of the trunk spinal cord, with dorsal up. (A,A’) At 53 hpf, dorsally migrated *olig2*^+^ OPCs are *nkx2.2a^+^* (filled arrowheads), indicating oligodendrocyte fate. (B,B’) At 60 hpf, some dorsally migrated *olig2*^+^ OPCs are *nkx2.2a*^*−*^ (outlined arrowheads). (D) Schematic representation of photoconversion experiment and result. Exposure of *Tg(olig2:Kaede);Tg(nkx2.2a:EGFP-CaaX)* to uv wavelength light at 30 hpf converts all *olig2*:Kaede^+^ cells from green to red. Motor neurons remain red because they no longer transcribe *olig2* whereas OPCs continue to express *olig2*, forming newly synthesized green Kaede. OPCs that initiated *olig2* expression prior to photoconversion are yellow whereas those that initiate *olig2* expression after photoconversion are green. To determine subtype identity, Kaede was photoconverted again at 55 hpf, revealing *nkx2.2a*:EGFP-CaaX expression marking OPCs fated for oligodendrocyte development. (E,E’) Projection of confocal image stack of 55 hpf embryo that had been photoconverted at 55 hpf. OPCs that are both green and red (filled arrowheads) and only green (outlined arrowheads) are evident. (F,F’) Image of same embryo shown in D following second photoconversion at 55 hpf. OPCs that were green and red are *nkx2.2a*^*^+^*^, indicating oligodendrocyte fate, whereas OPCs that were only green are *nkx2.2a*^*−*^.

Does the sequential appearance of *nkx2.2a^+^* and *nkx2.2a^−^* OPCs reflect sequential specification of distinct progenitors? In an earlier study we showed that motor neurons and OPCs are produced by distinct neural progenitors that initiate *olig2* expression at different times in a process we called progenitor recruitment (Ravanelli and Appel, 2015). Here we undertook a similar strategy to determine if OPCs that give rise to oligodendrocytes or persist as OPCs also initiate *olig2* expression at different times. As a tool to test the timing of *olig2* expression, we used the transgenic line *Tg(olig2:Kaede)*, which expresses the photoconvertable Kaede protein (Zannino and Appel, 2009). Photoconversion of Kaede using ultraviolet light causes an irreversible conversion of Kaede^*green*^ to Kaede^*red*^. Cells that continue to express *olig2*:Kaede following photoconversion appear yellow (Kaede^*yellow*^), from the combination of new Kaede^*green*^ and photoconverted Kaede^*red*^ (Figure 2D). By contrast, cells that initiate *olig2*:Kaede expression only after photoconversion appear green. To help discriminate between oligodendrocytes and later-born OPCs, we also included *Tg(nkx2.2a:EGFP-CaaX)*. Our previous experiments had shown that most motor neurons of *Tg(olig2:Kaede)* embryos photoconverted at 24 hpf and imaged at 72 hpf were Kaede^*red*^ whereas all dorsally migrated *olig2*^+^ cells were Kaede^*green*^, indicating that cells that developed as motor neurons and glial cells initiated *olig2* expression sequentially (Ravanelli and Appel, 2015). By contrast, when we photoconverted at 30 hpf and imaged at 55 hpf, some dorsal cells were Kaede^*yellow*^ (Figure 2E,E’), indicating that these cells expressed Kaede both before and after photoconversion, whereas others were only Kaede^*green*^, revealing that they initiated Kaede expression after photoconversion. To determine the subtype identity of these cells we repeated the photoconversion and imaging at 55 hpf. 93% of Kaede^*yellow*^ cells (n=142 cells, 21 embryos) were *nkx2.2a:*EGFP-CaaX^+^ (Figure 2F,F’). These data indicate that cells fated to differentiate as *nkx2.2a*^+^ oligodendrocytes initiated *olig2* expression before cells that did not express *nkx2.2a*. We therefore conclude that, during development, zebrafish larvae produce two distinct subpopulations of spinal cord OPCs: one population that initiates *olig2* expression early and rapidly differentiates as oligodendrocytes and a second population that initiates *olig2* expression later and persists into larval stage as OPCs.

### Different OPC subpopulations are specified by distinct levels of Shh signaling

The morphogen Shh patterns the ventral neural tube in a concentration-dependent manner. In particular, distinct, dorsoventrally arrayed populations of progenitors express different transcription factors as a result of graded levels of Shh signaling activity (Dessaud et al., 2008). For example, high levels of Shh cause the ventral-most progenitors of the p3 domain to express Nkx2.2 whereas slightly lower levels of Shh induce Olig2 expression by progenitors of the adjacent, more dorsally positioned pMN domain (Dessaud et al., 2008). Subsequently, a temporal rise in Shh signaling activity causes Nkx2.2 expression to expand dorsally into the pMN domain coincident with the switch from motor neuron to OPC production (Agius et al., 2004; Fu et al., 2002; Oustah et al., 2014; Touahri et al., 2012; Zhou et al., 2001). Could the sequential formation of distinct OPC subpopulations by progenitors that initiate *olig2* expression at different times result from different levels of Shh signaling?

To test this possibility, we used pharmacological and transgenic expression approaches to modulate Shh signaling levels. To lower signaling, we used the Smoothened inhibitor cyclopamine. In earlier work we had established that cyclopamine applied at 5 and 10 μM concentrations beginning at 24 or 30 hpf completely blocked OPC formation (Park et al., 2004; Ravanelli and Appel, 2015). To determine if different concentrations of cyclopamine quantitatively reduces signaling strength, we treated embryos with 5.0, 0.5 and 0.05 μM cyclopamine from 24-48 hpf, the period during which OPCs initially form, and performed in situ RNA hybridization to detect transcripts encoded by the Hedgehog pathway target gene *ptch2*. Transverse sections through the trunk spinal cord of embryos treated and processed in parallel revealed an apparent dose-dependent reduction of *ptch2* transcripts (Figure 3A). Additionally, we measured transcripts encoded by *ptch2* and *gli1*, which similarly is regulated by Hedgehog signaling, using semi-quantitative RT-PCR. This also revealed reduction of *ptch2* and *gli1* transcripts, although the differences between control, 0.05 and 0.5 μM were small and did not reach statistical significance as defined by p<0.05 (Figure 3B).

**Figure 3.**
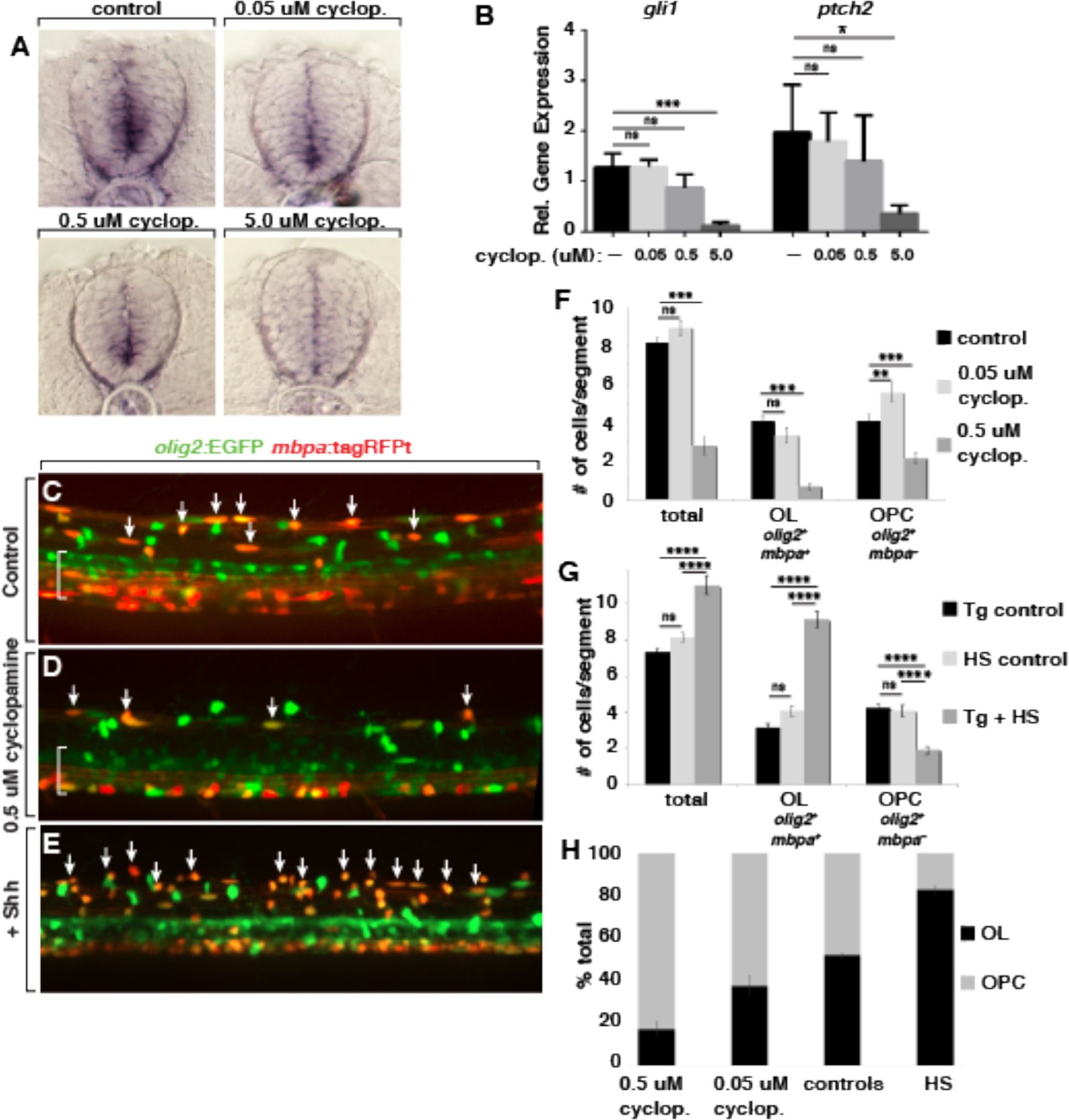
Different Levels of Shh Signaling Specify OPC and Oligodendrocyte Fates. (A) Representative images of transverse sections of trunk spinal cord obtained from 48 hpf embryos processed, in parallel, for in situ RNA hybridization to detect *ptch2* expression. Embryos were treated beginning at 24 hpf with control solution or 0.05, 0.5 or 5.0 μM cyclopamine (cyclop.). (B) Relative levels of *gli1* and *ptch2* transcripts detected by semi-quantitative RT-PCR. n=5 biological replicates with 20 larvae pooled per sample. (C-E) Representative images of confocal stacks of living 5 dpf *Tg(olig2:EGFP);Tg(mbpa:tagRFPt)* larvae at the level of the trunk spinal cord. Dorsal spinal cord is toward the top and brackets indicate the pMN domain in ventral spinal cord. Arrows mark *olig2^+^ mbpa^+^* oligodendrocytes. (C) Larva treated with control solution. (D) Larva treated with 0.5 μM cyclopamine. (E) Heat shocked larva carrying the *Tg(hsp70l:shha-EGFP)* transgene in addition to the reporter transgenes. (F) Average number of dorsal *olig2*^+^ cells, *olig2^+^ mbp^+^* oligodendrocytes (OL) and *olig2*^+^ *mbp*^*−*^ OPCs in control (n=50 embryos, 2676 cells), 0.5 μM cyclopamine-treated (n=17 embryos, 285 cells) and 0.05 μM cyclopamine-treated (n=7 embryos, 624 cells) larvae. (G) Average number of dorsal *olig2*^+^ cells, *olig2*^+^ *mbp*^+^ oligodendrocytes and *olig2*^+^ *mbp*^*−*^ OPCs in Tg control (heat-shocked larvae lacking *hsp70l:shha-EGFP* transgene; n=19 larvae, 842 cells), HS control (*hsp70l:shha-EGFP* larvae not subjected to heat-shock; n=10 larvae, 490 cells) and Tg ^+^ HS (heat-shocked *hsp70l:shha-EGFP* larvae; n=10 larvae, 663 cells). (H) Proportion of all dorsal *olig2*^+^ cells consisting of *olig2*^+^ *mbp*^+^ oligodendrocytes and *olig2*^+^ *mbp*^*−*^ OPCs from cyclopamine treatment and heat-shock experiments. * < 0.05, ** < 0.01, *** < 0.001, **** < 0.0001. ns, not significant, where p > 0.05. Data are presented as mean ± SEM. qPCR data evaluated using unpaired t test with Welsh correction. Cell count data evaluated using unpaired, two tailed t test.

We next tested if different levels of Hedgehog signaling specify neural progenitors for different OPC fates. We first assessed cell fates in living larvae by treating *Tg(olig2:EGFP);Tg(mbpa:tagRFPt)* embryos beginning at 24 hpf and counting cells that migrated into dorsal spinal cord. Whereas 5 μM cyclopamine completely blocked formation of all oligodendrocyte lineage cells (Ravanelli and Appel, 2015), 0.5 μM cyclopamine reduced the number of dorsal *olig2:*EGFP^+^ cells to about 30% of normal and 0.05 μM cyclopamine had no effect (Figure 3C,D,F). Notably, whereas 0.5 μM cyclopamine reduced the numbers of both oligodendrocytes (*olig2:*EGFP^+^ *mbp:*tagRFPt^+^) and OPCs (*olig2:*EGFP^+^ *mbp:*tagRFPt^−^), 0.05 μM cyclopamine did not change the number of oligodendrocytes but increased the number of OPCs (Figure 3F). Thus, formation of oligodendrocytes appears to require higher levels of Hedgehog signaling than does formation of OPCs that persist into larval stage. We therefore predicted that elevation of Hedgehog signaling would preferentially promote formation of oligodendrocytes. To test the effects of elevated Hedgehog signaling we used the transgenic line *Tg(hsp70l:shha-EGFP)*, which expresses Shh fused to EGFP under control of heat-responsive elements from the *hsp70l* gene (Shen et al., 2013). We heat shocked *Tg(hsp70l:shha-EGFP);Tg(olig2:EGFP);Tg(mbpa:tagRFPt)* embryos at 24 hpf and selected those with high levels of Shh-EGFP fluorescence for analysis. At 5 dpf, these larvae had fewer *olig2:*EGFP^+^ *mbpa:*tagRFPt^−^ OPCs but substantially more *olig2:*EGFP^+^ *mbpa:*tagRFPt^+^ oligodendrocytes than control larvae (Figure 3E,G), consistent with our prediction. To better assess the effects of different levels of Hedgehog signaling we graphed the ratios of oligodendrocytes and OPCs formed under each condition. This revealed an association between Shh signaling strength and cell fate whereby lower levels of Shh signaling favored formation of *olig2:*EGFP^+^ *mbpa:*tagRFPt^−^ OPCs and higher levels preferentially promoted formation of *olig2:*EGFP^+^ *mbpa:*tagRFPt^+^ oligodendrocytes (Figure 3H).

To test the validity of these results, we treated *Tg(cspg4:mCherry);Tg(mbpa:EGFP)* embryos with cyclopamine and examined expression of oligodendrocyte and OPC markers in tissue sections obtained from the trunk spinal cords of 5 dpf larvae. Neither 0.05 μM or 0.5 μM cyclopamine changed the number of OPCs significantly but decreased the number of oligodendrocytes in a dose-responsive manner (Figure 4A-D). Graphing the ratio of oligodendrocytes to OPCs indicated, similarly to the previous experiment, that reducing Hedgehog signaling increased the proportion of OPCs to oligodendrocytes (Figure 4E). We also performed the same treatments using *Tg(cspg4:mCherry)* embryos and *Tg(mbpa:EGFP)* embryos, which we then processed at 5 dpf to detect Sox10 using immunohistochemistry and analyzed in transverse tissue sections. As with the previous experiment, both marker combinations revealed that cyclopamine reduced the number of oligodendrocytes in a dose-responsive manner (Figure 4F,G). Additionally, whereas 0.5 μM slightly reduced the number of OPCs, larvae treated with 0.05 μM cyclopamine had more OPCs than controls (Figure 4F,G). Taken all together, these data indicate that spinal cord oligodendrocytes and larval OPCs are specified by distinct levels of Hedgehog signaling, whereby relatively higher levels promote oligodendrocyte formation and lower levels favor production of OPCs that persist into larval stage.

**Figure 4.**
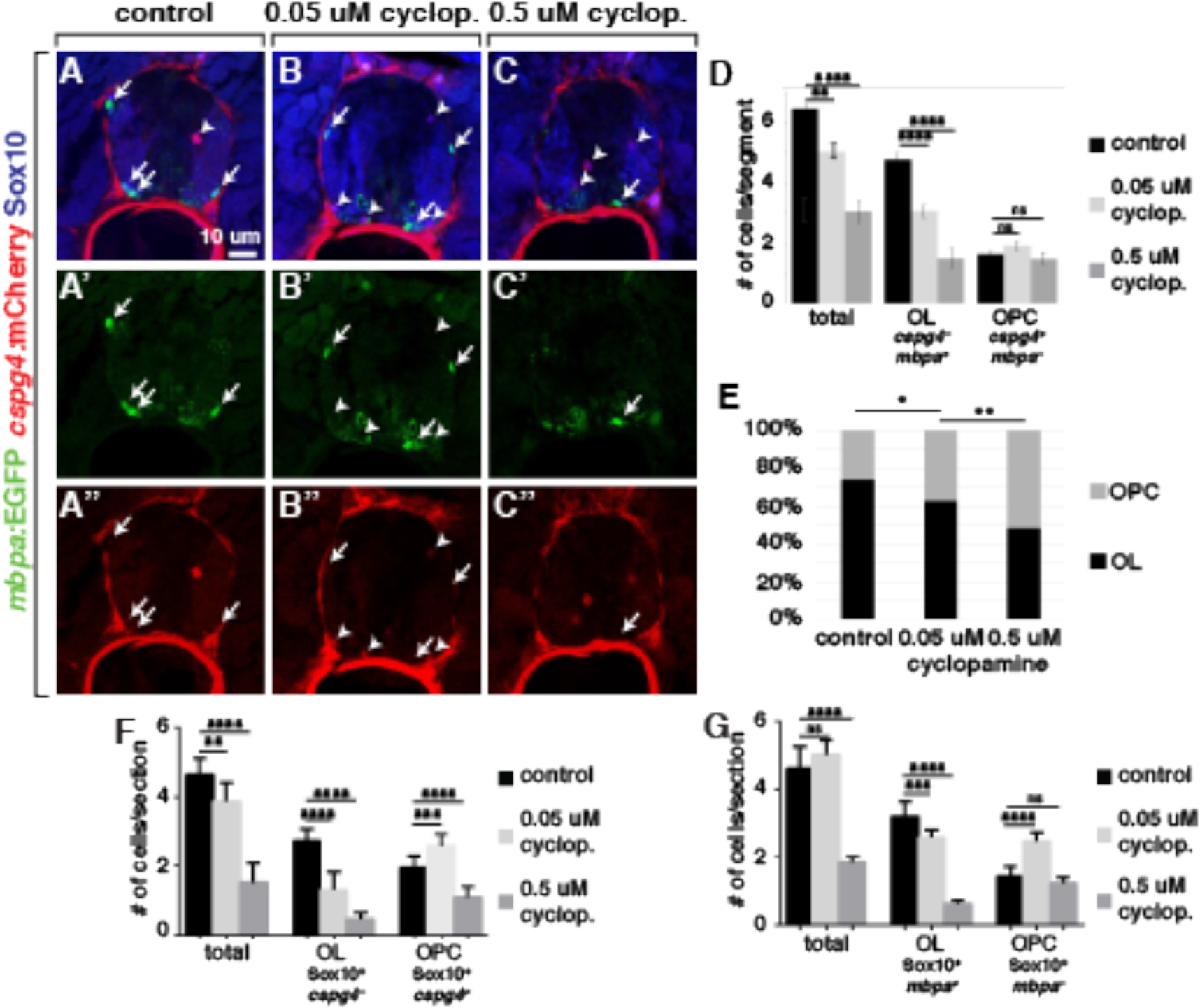
Changing the level of Hedgehog signaling changes the proportions of spinal cord oligodendrocytes and OPCs. (A-C”) Representative images of transverse sections through the trunk spinal cords of 5 dpf *Tg(cspg4:mCherry);Tg(mbpa:EGFP)* larvae and processed to detect Sox10 expression (blue). Arrows indicate *mbpa:*EGFP^+^ oligodendrocytes and arrowheads mark *cspg4:*mCherry^+^ OPCs. (D) Average number of total *cspg4*^+^ and *mbpa*^+^ cells, *cspg4*^+^ *mbp*^−^ OPCs and *cspg4*^−^ *mbp*^+^ oligodendrocytes in control (n=10 larvae, 643 cells), 0.5 μM cyclopamine-treated (n=10 larvae, 307 cells) and 0.05 μM cyclopamine-treated (n=10 larvae, 502 cells). (E) Proportions of *cspg4*^+^ *mbp*^−^ OPCs and *cspg4*^−^ *mbp*^+^ oligodendrocytes. (F) Average number of total Sox10^+^ *cspg4*^+^> and Sox10^+^ *cspg4^−^* cells, Sox10^+^ *cspg4^+^* OPCs and Sox10^+^ *cspg4^−^* oligodendrocytes in control (n=10 larvae), 0.5 μM cyclopamine-treated (n=10 larvae) and 0.05 μM cyclopamine-treated (n=10 larvae). p values are indicated by asterisks. * < 0.05, ** < 0.01, *** < 0.001, **** < 0.0001. ns, not significant, where p > 0.05. Data are presented as mean ± SEM. Cell count data evaluated using unpaired, two tailed t test.

### Notch signaling promotes oligodendrocyte specification by enhancing Hedgehog signaling activity

We previously showed that Notch signaling is necessary for specification of OPCs and that elevated Notch activity can promote the formation of excess oligodendrocyte lineage cells (Park and Appel, 2003; Snyder et al., 2012). However, the mechanism by which Notch signaling promotes OPC specification has not been known. Because other studies have indicated that Notch signaling can increase the strength of Hedgehog signal transduction (Huang et al., 2012; Kong et al., 2015) we tested the possibility that Notch regulates OPC specification by regulating neural progenitor responsiveness to Hedgehog ligands. To reduce Notch signaling, we used the gamma-secretase inhibitor LY411575, which effectively inhibits cleavage of the intracellular domain of Notch receptors thereby blocking Notch signaling activity. We first tested the effect of Notch inhibition on Hedgehog signaling by treating *Tg(ptch2:Kaede)* embryos, which express Kaede fluorescent protein under control of *ptch2* regulatory DNA (Huang et al., 2012). Treatment with 2.5 μM LY411575 from 24-48 hpf reduced Kaede fluorescence about 30% relative to control embryos (Figure 5A,B,D). For comparison, treatment with 5 μM cyclopamine for the same period reduced Kaede fluorescence approximately 70% (Figure 5C,D). By contrast, 5 μM cyclopamine did not change the fluorescence produced by a *Tg(her4.3:dRFP)* transgene (Figure 5F-H), which serves as a Notch reporter (Yeo et al., 2007), indicating that Hedgehog signaling does not influence Notch signaling activity.

**Figure 5.**
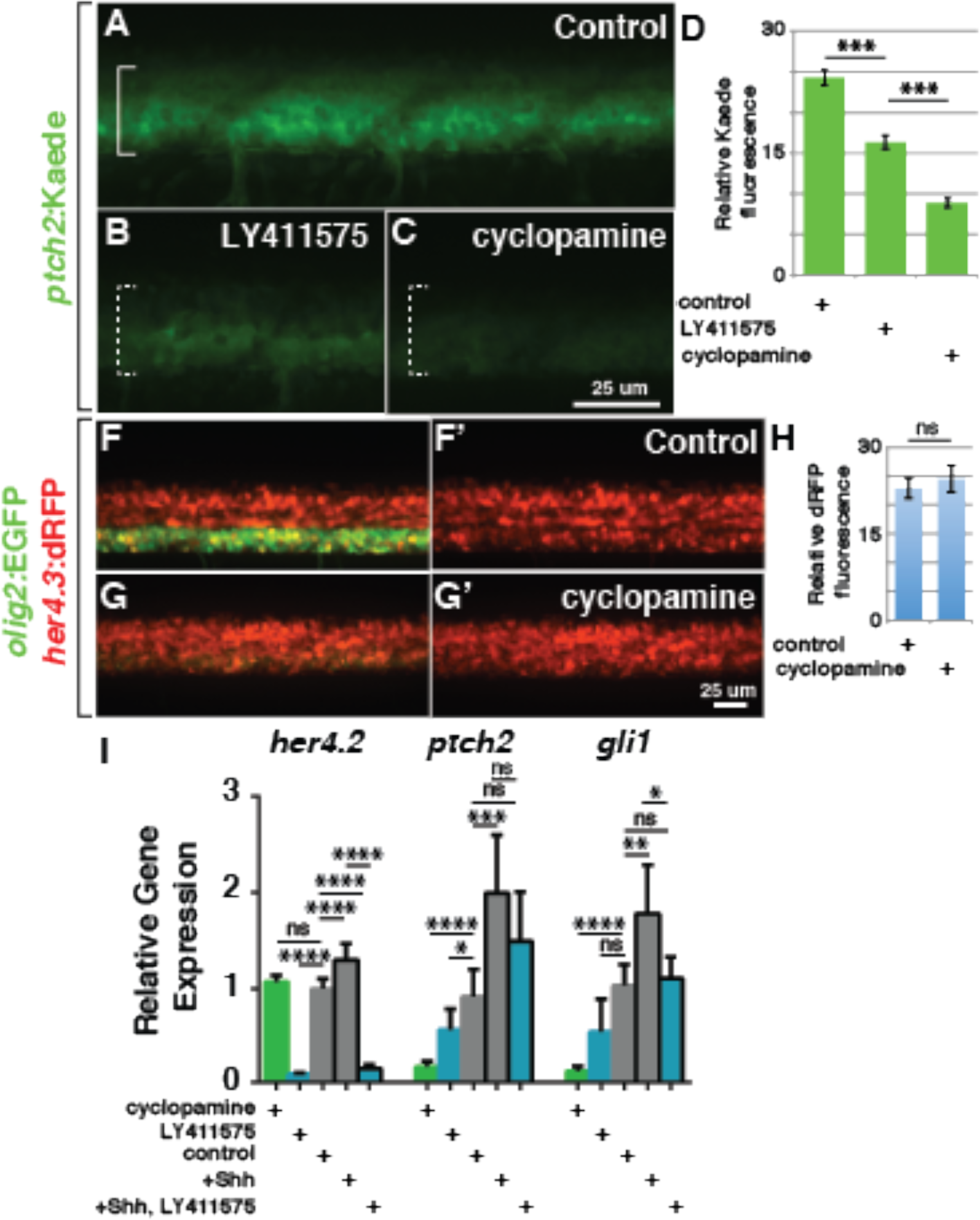
Notch Activity Enhances Shh Signaling. (A-C) Representative confocal images of living 48 hpf *Tg(ptch2:Kaede)* embryos at the level of the trunk spinal cord with dorsal up. Brackets mark ventral spinal cord. (A) Embryo treated with control solution. (B) Embryo treated with 2.5 μM LY411575 to inhibit Notch signaling. (C) Embryo treated with 5 μM cyclopamine. (D) Relative Kaede fluorescence intensity measurements for control (n=5), LY411575 treated (n=6) and cyclopamine treated (n=5) embryos. (F-G’) Representative confocal images of living 48 hpf *Tg(olig2:EGFP);Tg(her4.3:dRFP)* embryos at the level of the trunk spinal cord. (F) Embryo treated with control solution. (G) Embryo treated with 5 μM cyclopamine. (H) Relative dRFP fluorescence intensity measurements for control (n=18) and cyclopamine-treated (n=11) *Tg(olig2:EGFP);Tg(her4.3:dRFP)* embryos. (I) Relative levels of *her4.2, ptch2* and *gli1* transcripts. Cyclopamine and LY411575 concentrations were 5.0 and 2.5 μM, respectively. ^+^Shh indicates heat shocked *Tg(hsp70l:shha-EGFP)* embryos. p values are indicated by asterisks. * < 0.05, ** < 0.01, *** < 0.001, **** < 0.0001. ns, not significant, where p > 0.05. Data are presented as mean ± SEM. Fluorescence intensity data evaluated using two-tailed t test. qPCR data evaluated using unpaired t test with Welsh correction.

To validate and extend these results we performed a series of semi-quantitative RT-PCR experiments to measure levels of transcripts encoded by Hedgehog and Notch pathway target genes. Consistent with the quantitative imaging data, cyclopamine treatment from 24-48 hpf significantly reduced *ptch2* and *gli1* but not *her4* transcript levels (Figure 5I). Treatment with 2.5 μM LY411575 nearly eliminated *her4.2* transcripts and reduced both *ptch2* and *gli1* transcript levels although the difference in *gli1* levels was not statistically significant relative to control (Figure 5I). Furthermore, whereas heat shock induction of Shh-EGFP expression elevated *ptch2* and *gli1*, but not *her4* RNA levels, simultaneous treatment of heat shocked embryos with LY411575 reduced the amount of all three transcripts, although the reduction of *ptch2* transcripts did not reach statistical significance as determined by p<0.05. (Figure 5I). These data are consistent with the idea that Notch activity enhances Shh pathway signaling strength.

To test the hypothesis that Notch signaling is required for the specification of myelin-fated OPCs, we exposed embryos to the Notch inhibitor from 24-72 hpf, the period of progenitor recruitment and OPC specification. We had previously shown that strong inactivation of Notch signaling prior to OPC specification using the Notch inhibitor DAPT or heat-induced expression of a dominant-negative form of RBP Jκ entirely blocked OPC formation (Kim et al., 2008; Park et al., 2005b; Snyder et al., 2012). Here, as with our cyclopamine experiments, we tested different concentrations of Notch inhibitor to learn if different levels of Notch signaling influence formation of distinct OPC subpopulations. Compared to the control (Figure 6A), 1 μM LY411575 prevented formation of nearly all dorsally migrated *olig2:*EGFP^+^ cells (Figure 6D) whereas 0.5 μM LY411575 reduced the number of *olig2:*EGFP^+^ *mbpa*:mCherry^+^ oligodendrocytes without changing the number of *olig2:*EGFP^+^ *mbpa*:mCherry^−^ cells (Figure 6B,D). Graphing the data as a proportion of the entire cell population revealed that, like our Shh inhibitor experiments, reducing Notch signaling resulted in fewer oligodendrocytes relative to OPCs (Figure 6E). We interpret these results to mean that formation of oligodendrocytes requires higher levels of Notch signaling than formation of OPCs that persist into larval stage.

**Figure 6.**
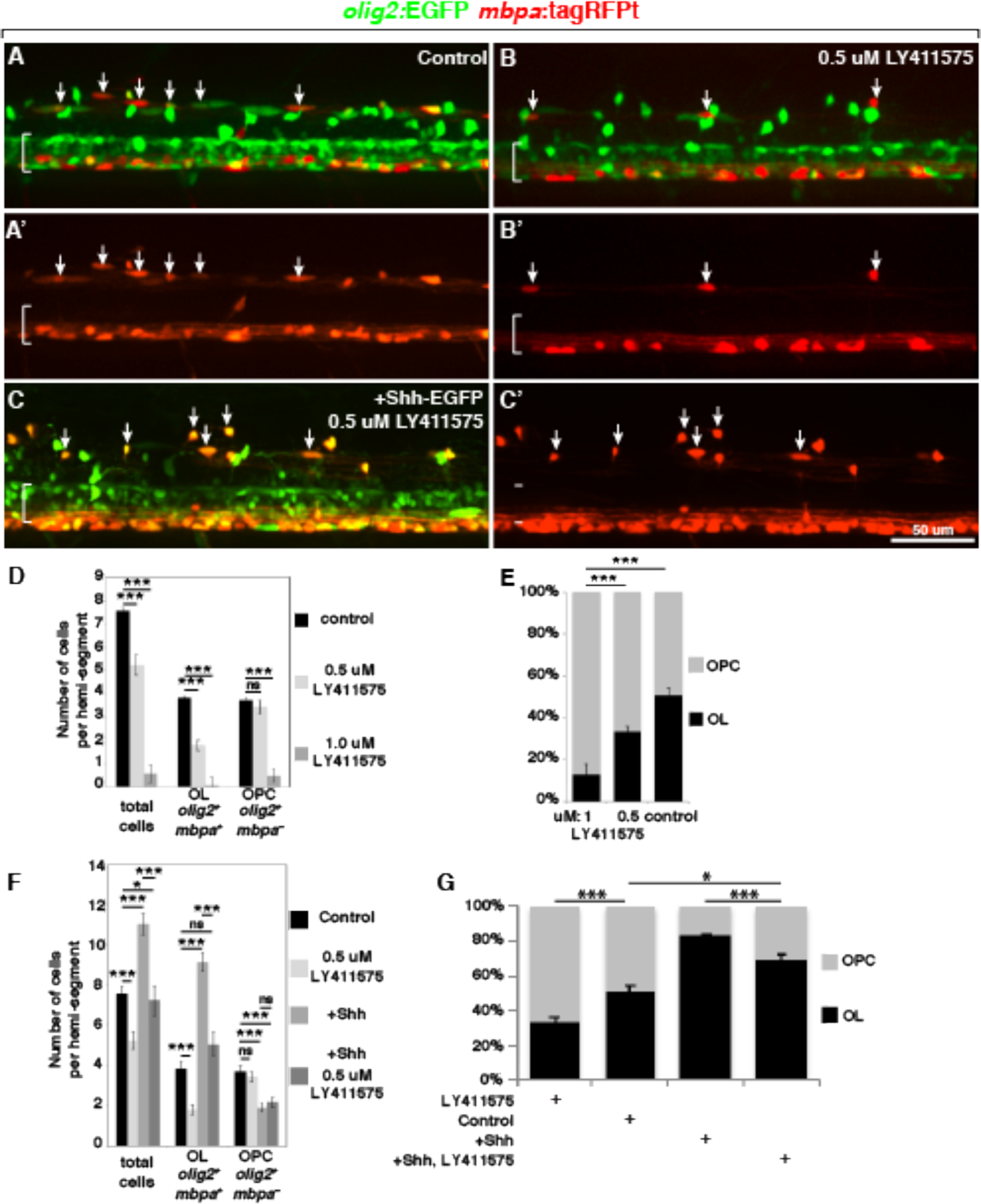
Notch Activity Modulates Shh Signaling to Specify Oligodendrocytes and OPCs. (A-C’) Representative projections of confocal stack images of living 5 dpf *Tg(olig2:EGFP);Tg(mbpa:tagRFPt)* larvae. Dorsal is up and brackets mark the pMN domain of ventral spinal cord. Arrows indicate *olig2*^+^ *mbpa*^+^ oligodendrocytes. Panel pairs show *olig2*:EGFP and *mbpa:*tagRFPt expression combined and *mbpa:*tagRFPt expression alone. (A,A’) Larva treated with control solution. (B,B’) Larvae treated with 0.5 μM LY411575. (C,C’) Heat shocked larva carrying the *Tg(hsp70l:shha-EGFP)* transgene in addition to the reporter transgenes and treated with 0.5 μM LY411575. (D) Average number of total dorsal spinal cord *olig2*^+^ cells, *olig2*^+^ *mbp*^+^ oligodendrocytes and *olig2*^+^ *mbpa*^−^ OPCs in control (n=10, 454 cells), 0.5 μM LY411575 treated (n=11, 346 cells) and 1.0 μM LY411575 treated (n=17, 60 cells) larvae. (E) The effect of Notch inhibition on the proportions of *olig2*^+^ *mbpa*^+^ oligodendrocytes and *olig2*^+^ *mbpa*^−^ OPCs. (F) Average number of total number of dorsal spinal cord *olig2*^+^ cells, *olig2*^+^ *mbpa*^+^ oligodendrocytes and *olig2*^+^ *mbpa*^−^ OPCs in control, 0.5 μM LY411575 treated, Shh overexpressing and Shh overexpressing ^+^ 0.5 μM LY411575 treated (n=21, 760 cells) larvae. For this graph the control data were combined from Figures 3F and 5D. The LY411575 data are from Figure 5D and the ^+^Shh data are from Figure 3F, shown here for comparison. (G) Proportion of all dorsal *olig2*^+^ cells consisting of *olig2*^+^ *mbpa*^+^ oligodendrocytes and *olig2*^+^ *mbpa*^−^ OPCs following Notch inhibition and Shh overexpression in the presence and absence of Notch inhibitor. p values are indicated by asterisks. * < 0.05, ** < 0.01, *** < 0.001, **** < 0.0001. ns, not significant, where p > 0.05. Data are presented as mean ± SEM. Data were evaluated using unpaired, two-tailed t test.

Does the ability of elevated Shh levels to drive formation of excess oligodendrocytes require Notch function? Indeed, heat-shocked *Tg(hsp70l:shha-EGFP)* embryos treated with 0.5 μM LY411575 had approximately normal numbers of oligodendrocytes relative to control (Figure 6C,C’,F) and proportionally fewer oligodendrocytes than heat shocked *Tg(hsp70l:shha-EGFP)* embryos without Notch inhibitor (Figure 6G). Thus, Notch activity appears to be necessary for neural cells to respond to elevated levels of Shh, consistent with the idea that Notch activity modulates Shh signaling strength.

### Small changes in Shh and Notch signaling do not dramatically alter pMN progenitor domain refinement

Several lines of evidence have shown that the expression domain of *Nkx2.2/nkx2.2a* expands dorsally into the pMN domain concomitant with OPC specification (Fu et al., 2002; Kucenas et al., 2008b; Oustah et al., 2014; Touahri et al., 2012; Zhou et al., 2001). Additionally, treatment of zebrafish embryos with 1 μM cyclopamine from 30-48 hpf blocked expansion of the *nkx2.2a* expression domain and resulted in a deficit of *olig2*:EGFP^+^ OPCs (Oustah et al., 2014). Thus, persistent, high-level Shh signaling appears to drive new gene expression in pMN progenitors to specify OPC formation following motor neuron production. We therefore investigated *nkx2.2a* expression following our own cyclopamine and LY411575 treatment conditions (Figure 7A-F). To quantify the effects of these treatments, we measured the height of the *nkx2.2a* expression domains and graphed them as a proportion of the total height of the neural tube. In vehicle-treated control embryos, the height of the *nkx2.2a* expression domain was about 30% of the controls (Figure 7G,H). The height of the *nkx2.2a* expression domain was slightly reduced in embryos treated with 0.05 μM cyclopamine, although it was not statistically different in those treated with 0.5 μM cyclopamine (Figure 7G). Treatment with the Notch inhibitor had a stronger effect, with the height of the *nkx2.2a* expression domain reduced in embryos treated with either 0.5 or 1.0 μM LY411575 (Figure 7H). These data suggest that our Shh and Notch inhibitor conditions can alter the allocation of progenitors that develop as oligodendrocytes or persist as OPCs without dramatically affecting *nkx2.2a* expression dynamics.

**Figure 7.**
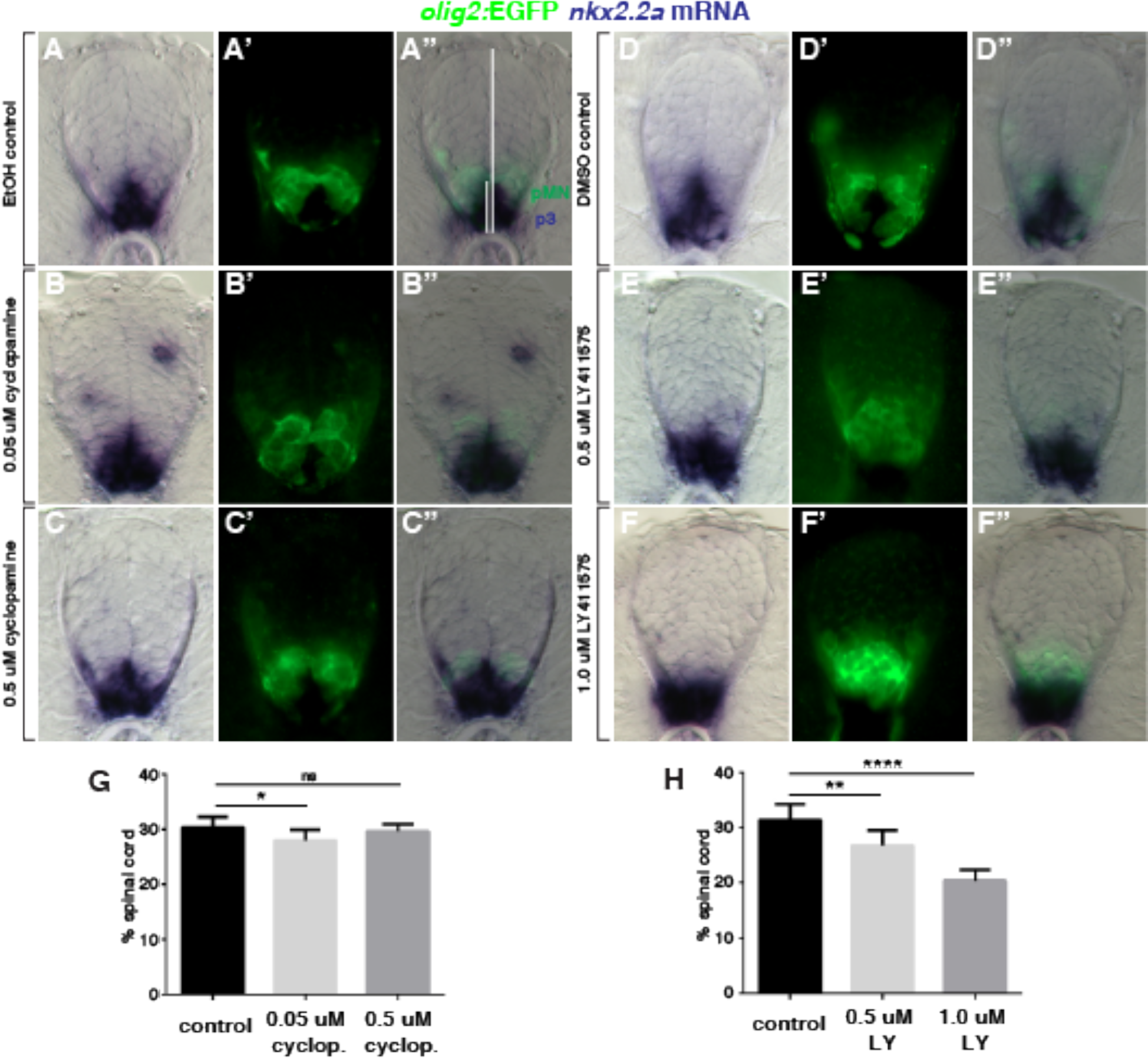
Effects of Shh and Notch inhibitors on *nkx2.2a* expression dynamics. (A-F) Representative examples of differential interference contrast (DIC), epifluorescent and combined images of transverse sections of *Tg(olig2:EGFP)* embryos treated from 24-48 hpf with control solutions (A,D), cyclopamine (B,C) or LY411575 (E,F) and processed to detect *nkx2.2a* mRNA expression. *nkx2.2a* expression marks the p3 domain whereas *olig2*:EGFP expression marks the pMN domain. White lines on panel A” indicate height measurements for the *nkx2.2a* expression domain and spinal cord. Averaged percent height of the spinal cord occupied by *nkx2.2a* expression for EtOH treated control (n=8), 0.05 μM cyclopamine treated (n=7) and 0.5 μM cyclopamine treated (n=7) embryos. (H) Averaged percent height of the spinal cord occupied by *nkx2.2a* expression for DMSO treated control (n=8), 0.5 μM LY411575 treated (n=8) or 0.5 μM LY411575 treated (n=8) embryos. p values are indicated by asterisks. * < 0.05, ** < 0.01, **** < 0.0001. ns, not significant, where p > 0.05. Data are presented as mean ± SEM. Data were evaluated using unpaired, two-tailed t test.

### OPCs differentiate as Mbp^+^ oligodendrocytes independently of Shh and Notch signaling

We interpret the above data to mean that Shh and Notch signaling regulate the number of neural progenitors that are specified as OPCs fated for oligodendrocyte development. An alternative possibility is that Shh and Notch act later to regulate OPC proliferation and differentiation as oligodendrocytes. To test this, we repeated the inhibitor and overexpression experiments after OPCs were already formed.

*Tg(olig2:EGFP);Tg(mbpa:tagRFPt)* larvae treated with 10 μM cyclopamine from 72 hpf to 5 dpf had no statistically significant differences in the number of dorsal spinal cord *olig2:*EGFP^+^ cells compared to control larvae (Figure 8A,B,D) Similarly, 5 dpf *Tg(olig2:EGFP);Tg(mbpa:tagRFPt);Tg(hsp70l:shha-EGFP)* larvae that had been heat shocked at 72 hpf had no statistically significant differences in the number of dorsal spinal cord *olig2:*EGFP^+^ cells compared to control larvae (Figure 8C,E). Treatment with high doses of Notch inhibitor (10 μM or 25 μM) did not prevent the differentiation of OPCs into *mbpa:*tagRFPt^+^ oligodendrocytes by 5 dpf relative to controls (Figure 8F-H) and did not alter the ratio between oligodendrocytes and *olig2:*EGFP^+^ *mbpa*:tagRFPt^−^ OPCs at 5 dpf (Figure 8I). These data support the idea that Shh and Notch signaling function during progenitor specification to regulate formation of OPCs that differentiate as oligodendrocytes, rather than during their subsequent differentiation.

**Figure 8.**
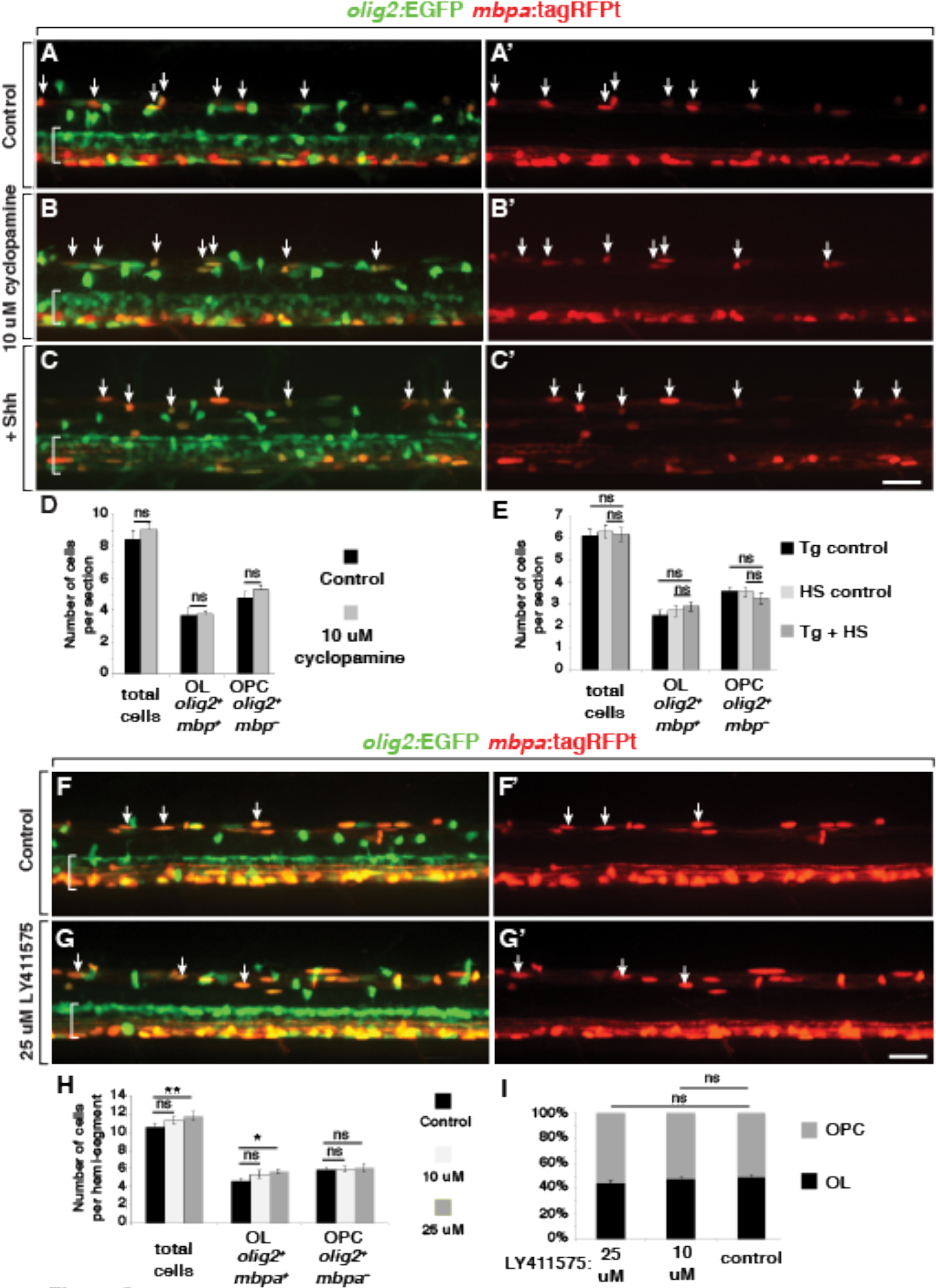
Manipulation of Shh and Notch Signaling Following Specification Does Not Alter Oligodendrocyte and OPC Formation. (A-C’, F-G’) Representative projections of confocal stack images of living 5 dpf *Tg(olig2:EGFP);Tg(mbpa:tagRFPt)* larvae. Dorsal is up and brackets mark the pMN domain of ventral spinal cord. Arrows indicate *olig2*^+^ *mbpa*^+^ oligodendrocytes. Panel pairs show *olig2*:EGFP and *mbpa:*tagRFPt expression combined and *mbpa:*tagRFPt expression alone. (A,A’) Larva treated with control solution. (B,B’) Larva treated with 10 μM cyclopamine beginning at 72 hpf. (C,C’) *hsp70l:Shh-GFP; olig2:EGFP; mbpa:tagRFPt* larva heat-shocked at 72hpf. (D) Average number of dorsally migrated *olig2*^+^ *mbpa*^+^ oligodendrocytes and *olig2*^+^ *mbpa*^−^ OPCs for control (n=24 larvae,1042 cells) and 10 μM cyclopamine (n=11 larvae, 596 cells) treatments. (E) Average number of dorsally migrated *olig2*^+^ *mbpa*^+^ oligodendrocytes and *olig2*^+^ *mbpa*^−^ OPCs for transgenic (Tg) control (n=8 larvae), heat-shock (HS) control (n=5 larvae) and ^+^Shh (n=16 larvae, 590 cells) treatments. (F,F’) Larva treated with control solution. (G,G’) Larva treated with 25 μM Notch inhibitor LY411575 beginning at 72 hpf. (H) Average number of dorsally migrated *olig2*^+^ *mbpa*^+^ oligodendrocytes and *olig2*^+^ *mbpa*^−^ OPCs in control (n=9 larvae, 571 cells),10 μM LY411575 (n=15 larvae, 1021 cells), and 25 μM LY411575 (n=15 larvae, 1064 cells) treatments. p values are indicated by asterisks. * < 0.05, ** < 0.01. ns, not significant, where p > 0.05. Data are presented as mean ± SEM. Data were evaluated using unpaired, two-tailed t test. All scale bars represent 25 um.

## DISCUSSION

OPCs divide and migrate to generate sufficient numbers of oligodendrocytes properly positioned for myelination. However, many OPCs persist into adulthood without differentiating as oligodendrocytes. Why do some OPCs produce oligodendrocytes whereas others do not?

One model of oligodendrocyte development is that oligodendrocyte lineage cells progress along a common, linear pathway of differentiation. In this model, OPCs that persist without becoming oligodendrocytes represent an early step in the pathway, with the implication that their differentiation toward myelinating cells is arrested. Consistent with this possibility, numerous signals have been identified that act as brakes on oligodendrocyte differentiation. For example, Wnts appear to be negative regulators of oligodendrocyte differentiation because expression of a constitutively active form of β-catenin in oligodendrocyte lineage cells reduced myelination (Fancy et al., 2009) whereas an antagonist of Wnt receptors stimulated formation of immature oligodendrocytes (Shimizu et al., 2005). LINGO-1, a transmembrane protein expressed by oligodendrocytes, inhibits myelination, potentially by homophilic interaction with LINGO-1 on axons (Jepson et al., 2012; Mi et al., 2005). The G-protein-coupled receptor GPR17 and the Notch receptor also negatively regulate oligodendrocyte myelination (Chen et al., 2009; Wang et al., 1998). Thus, signals that inhibit myelination could be primary determinants of OPC fate.

An alternative possibility is that distinct subsets of OPCs are specified for different fates. Supporting this possibility, a deficit of prefrontal cortex OPCs produced depressive-like behaviors in mice (Birey et al., 2015), suggesting that some OPCs have functions other than myelination. With this study we present evidence for two distinct OPC subpopulations in the spinal cord of zebrafish larvae. First, although both subpopulations express *olig2*, they initiate expression at different times, with progenitors developing as oligodendrocytes expressing *olig2* before those that persist as OPCs into larval stage. This is similar to our previous work that showed that progenitors that produce motor neurons initiate *olig2* before those that generate OPCs in a process we called progenitor recruitment (Ravanelli and Appel, 2015). We now extend the progenitor recruitment model to propose that motor neurons, rapidly myelinating oligodendrocytes and persistent OPCs arise from distinct spinal cord progenitors that sequentially initiate *olig2* expression. Having different, rather than common, progenitor origins is consistent with a view that these represent distinct cell types rather than cells at different stages along a common oligodendrocyte differentiation pathway. Second, small changes in Shh and Notch signaling have different effects on the formation of oligodendrocytes and OPCs, suggesting that they are specified by different levels of signaling. In particular, slightly higher levels of Shh and Notch signaling promote formation of OPCs that rapidly differentiate as oligodendrocytes. In the context of the progenitor recruitment model, we propose that an initial wave of progenitors that enter the pMN domain following motor neuron formation move more ventrally than following progenitors. Consequently, progenitors of the early wave are exposed to slightly higher levels of Shh, initiate expression of *nkx2.2a* and develop as oligodendrocytes whereas the following progenitors receive less Shh, do not express *nkx2.2a* and remain as OPCs. This finding has potentially important implications for methods to produce oligodendrocytes in vitro because slight differences in culture conditions could reduce OPC myelinating potential.

Previous studies have shown that Shh signaling within the developing neural tube is dynamic. For example, the response of neural cells to Shh exposure changes over time, in a process called temporal adaptation (Dessaud et al., 2007). Neighboring progenitors having different levels of Shh signaling can sort out to create homogeneous progenitor domains (Xiong et al., 2013). The source of Shh shifts over time from notochord, mesoderm underlying the neural tube, to floorplate, the most ventral cell type of the neural tube, and then to include cells of the adjacent p3 domain (Charrier et al., 2002; Oustah et al., 2014; Park et al., 2004). Finally, neuroepithelial cells move ventrally as motor neurons differentiate, changing their positions within the Shh signaling gradient (Ravanelli and Appel, 2015). Altogether, these studies indicate that changes in the Shh signaling gradient and responding cells during development can contribute to neural cell fate diversification.

A long-standing problem in vertebrate neural development is how apparently common populations of progenitors produce different kinds of neurons and glia. Current models frequently invoke asymmetric cell division and distribution of specification factors as mechanisms for generating different types of cells from common progenitors (Delaunay et al., 2017; Taverna et al., 2014). However, neurons and glia appear often to arise from distinct, instead of common, progenitors because most clonally related cell lineages in rodents consist only of neurons or glia (Grove et al., 1993; Luskin et al., 1988, 1993; McCarthy et al., 2001). How progenitors become restricted to particular neuronal and glial fates is not well understood. Together with our earlier work (Ravanelli and Appel, 2015), our current results raise the possibility that the movements of progenitors within gradients of signaling molecules can alter their signaling responses and diversify cell fate.

Although this study provides strong evidence that small differences in Hedgehog and Notch signaling can influence the development of OPCs that have distinct fates in larval zebrafish, several important questions remain unanswered. First, what is the origin and relationship of the neural progenitors that give rise to OPCs that differentiate as oligodendrocytes in early larval stages and those that remain as OPCs? Our earlier work showed that neuroepithelial cells move ventrally to enter the pMN domain just prior to OPC specification (Ravanelli and Appel, 2015). However, we currently lack transgenic reporters that identify these progenitors before they initiate *olig2* expression, leaving us with inexact knowledge of their identities prior to entering the pMN domain. Second, how do small differences in Hedgehog signaling specify distinct OPC subtypes? In zebrafish, the earliest known molecular distinction between OPC subtypes is *nkx2.2a* transcription. Determining the mechanisms that control *nkx2.2a* transcription in response to graded Hedgehog signaling should be an important step toward understanding OPC subtype specification. Third, what are the functions and fates of OPCs that do not differentiate as oligodendrocytes? Use of CRISPR/Cas9-mediated genome modification to create better tools to manipulate and permanently mark OPCs in zebrafish should provide a way forward to understand how they contribute to brain function.

## ACKNOWLEDGMENTS

We gratefully acknowledge Ajay Chitnis, Robert Kelsh, Rolf Karlstrom, Hae-Chul Park, Cheol-Hee Kim, Phil Ingham, Kristen Kwan, Chi-Bin Chien and Stefan Schulte-Merker for gifts of zebrafish lines and plasmids. DR274 and MLM3613 were gifts from Keith Joung (Addgene plasmids #42250 and #42251). This work was supported by the National Institutes of Health (NS40660 to B.A., CA08208613 to A.M.R., MN015442 to J.H.H), the National Multiple Sclerosis Society (FG 2024-A-1 to J.H.H.) and the Gates Frontiers Fund (gift to B.A.). The University of Colorado Anschutz Medical Campus Zebrafish Core Facility was supported by National Institutes of Health grant P30 NS048154. The authors declare that they have no financial or competing interests.

## AUTHOR CONTRIBUTIONS

Conceptualization, A.M.R and B.A.; Methodology, A.M.R.; Resources, A.M.R., Y.W. and J.H.H.; Investigation, A.M.R., C.A.K., R.K.P., M.J.D. and B.A.; Writing - Original Draft, A.M.R and B.A.; Writing’ Review & Editing, B.A., A.M.R. and C.A.K.; Funding Acquisition, B.A., A.M.R. and J.H.H.; Supervision, B.A.

